# Mu opioid receptor expression by nucleus accumbens inhibitory interneurons promotes affiliative social behavior

**DOI:** 10.1101/2024.10.28.620729

**Authors:** Carlee Toddes, Jessica L. Swanson, Emilia M. Lefevre, Cassandra L. Retzlaff, Lydia B. Cunitz, Ursula C. H. Girdwood, Lauryn Zugschwert, Sameer Khan, Erin Myhre, Elysia A. Gauthier, Ezequiel Marron Fernandez de Velasco, Brigitte L. Kieffer, Patrick E. Rothwell

**Affiliations:** Graduate Program in Neuroscience, University of Minnesota, Minneapolis, MN; Department of Neuroscience, University of Minnesota, Minneapolis, MN; Neuroscience Program & Department of Biology, University of St. Thomas, St. Paul, MN; Department of Pharmacology, University of Minnesota, Minneapolis, MN; INSERM U1329, University of Strasbourg Institute for Advanced Study, Strasbourg 67084, France

## Abstract

Mu opioid receptors in the nucleus accumbens regulate motivated behavior, including pursuit of natural rewards like social interaction as well as sensitivity to exogenous opioids. However, the specific circuit elements where mu opioid receptor expression promotes these motivated behaviors remain unclear. In mice of both sexes, we used a suite of genetic and viral strategies to conditionally delete mu opioid receptor expression from major neuron types in the nucleus accumbens, as well as monosynaptic inputs to this brain region. We pinpointed local inhibitory interneurons as an essential site of mu opioid receptor expression for typical social behavior. Mu opioid receptor deletion from inhibitory interneurons decreased both social interaction and preference, while shifting the ratio between synaptic excitation and inhibition onto medium spiny projection neurons, but did not alter exogenous opioid sensitivity. Conversely, mu opioid receptor deletion from medium spiny neurons altered exogenous opioid sensitivity but not social interaction, revealing a double-dissociation in the behavioral functions of mu opioid receptors expressed by nucleus accumbens projection neurons versus interneurons. Chemogenetic activation of inhibitory interneurons in the nucleus accumbens also decreased social interaction, mimicking the effect of mu opioid receptor deletion from these cells. Our findings suggest that circuit-specific engagement of mu opioid receptors could circumvent the risk of misuse associated with conventional exogenous opioids, with important implications for the pathophysiology and treatment of neuropsychiatric conditions involving social differences.

## INTRODUCTION

The mu opioid receptor (MOR) is expressed by cells in multiple reward-related brain circuits, including the striatum and the ventral tegmental area^1, 2^. MOR activation by exogenous opioids like morphine and fentanyl contributes to misuse of these drugs, and has been a major contribution to the opioid epidemic^3^. MOR activation in the nucleus accumbens (NAc) is sufficient to drive an array of motivated behaviors^2^, ranging from affiliative social interaction^4^ to food consumption^5^ to opioid self-administration^6^. The molecular circuitry underlying these behavioral changes can involve presynaptic mu opioid receptors expressed on axon terminals of synaptic inputs to the NAc^7, 8, 9^, as well as postsynaptic mu opioid receptors expressed by cells located within the NAc^10, 11, 12^. While constitutive mu opioid receptor knockout produces profound deficits in social behavior^13, 14, 15, 16^, the location and identity of key cell types where MOR expression promotes social behavior have not been resolved. This knowledge gap precludes a mechanistic comparison between the role of mu opioid receptors in social interaction versus other forms of motivated behavior. Such information is necessary to determine if enhancement of mu opioid receptor signaling to increase sociability carries the iatrogenic risk of misuse or abuse associated with conventional mu opioid receptor agonists.

MORs are expressed on virtually all NAc neuronal subtypes, including medium spiny neurons projection neurons that express dopamine receptors Drd1 (D1-MSNs) and Drd2 (D2-MSNs)^10^, as well as local interneurons^17^ (Fig. 1a). These interneurons include low-threshold spiking interneurons (LTSIs) marked by expression of somatostatin (Sst); fast-spiking interneurons (FSIs) marked by expression of parvalbumin; and cholinergic interneurons marked by expression of choline acetyltransferase^18^. In this study, we systematically knocked out Oprm1 (the gene encoding MOR) from the NAc in a cell type- and region-specific manner. In conditional Oprm1 knockout mice, pan-neuronal Oprm1 deletion resulted in impaired social behavior, as indicated by reduced social interaction. We found that deletion of Oprm1 from NAc MSNs affected sensitivity to exogenous opioid administration, but not social behavior. On the other hand, deletion of Oprm1 from NAc inhibitory interneurons impaired social behavior without affecting sensitivity to exogenous opioids. These data indicate that different MOR-expressing subpopulations in the NAc regulate different types of motivated behavior. This mechanistic understanding will aid in the development of novel, circuit-based therapeutic strategies to modify social behavior without abuse liability.

## RESULTS

To dissect the molecular circuitry through which mu opioid receptors regulate social behavior, we used mice with loxP sites flanking exons 2-3 of the Oprm1 gene^19^ (Oprm1^fl/fl^), which are excised in the presence of Cre recombinase (Fig. 1b). We introduced Cre using stereotaxic injection of adeno-associated viral vectors into the nucleus accumbens, while injecting separate control mice with an inactive virus. Prior to virus injection, we measured baseline social interaction with an age- and sex-matched conspecific (Fig. 1c). We then conducted a social interaction test following six weeks of virus expression, allowing us to compare behavior between groups for each social interaction session, as well as within groups across social interaction sessions. This design helps control for age-related changes in social behavior, as well as individual differences in overall sociability.

We first expressed Cre recombinase using AAV2retro, a viral serotype that is taken up by axon terminals and transported retrogradely^20^. Injection of this virus into the NAc allowed us to delete Oprm1 expression from presynaptic axon terminals as well as postsynaptic neurons (which have local axon collaterals), thus removing Oprm1 expression from all major NAc circuit elements (Fig. 1c). After measuring baseline social interaction, we performed bilateral injections of AAV2retro-hSyn-Cre-P2A-tdTomato into the NAc (Fig. 1d and Extended Data Fig. 1a). Control mice were injected with an inactive control virus (AAV2retro-hSyn-Flp-P2A-tdTomato), which expressed Flp recombinase instead of Cre recombinase. Statistical analysis across baseline and test sessions indicated a significant Session x Virus interaction (F_1,28_ = 4.43, p = 0.044, η_p_^2^ = 0.14), with significantly less social interaction in the Cre group versus controls during the test session (Fig. 1e). This pattern was also evident when a difference score (test – baseline) was compared between genotypes (Fig. 1f).

To determine if postsynaptic deletion of Oprm1 was sufficient to reproduce this phenotype, a separate cohort of Oprm1^fl/fl^ mice received bilateral injection of AAVdj-hSyn-Cre-GFP^21^ into the NAc (Extended Data Fig. 1b), to remove Oprm1 expression from neurons at the site of injection (Fig. 1g). Control mice were injected with virus expressing a catalytically inactive form of Cre (AAVdj-hSyn-deltaCre-GFP). Statistical analysis across baseline and test sessions (Fig. 1e) indicated a significant Session x Virus interaction (F_1,28_ = 6.41, p = 0.017, η_p_^2^ = 0.19), with significantly less social interaction in the Cre group at test versus baseline (Fig. 1h). This pattern was also evident when a difference score (test – baseline) was compared between genotypes (Fig. 1i). None of these viral manipulations affected behavioral sensitivity to acute morphine exposure (Extended Data Fig. 2). Together, these results indicate Oprm1 expression by postsynaptic NAc neurons is necessary to support normal levels of reciprocal social interaction.

**Figure 1.**
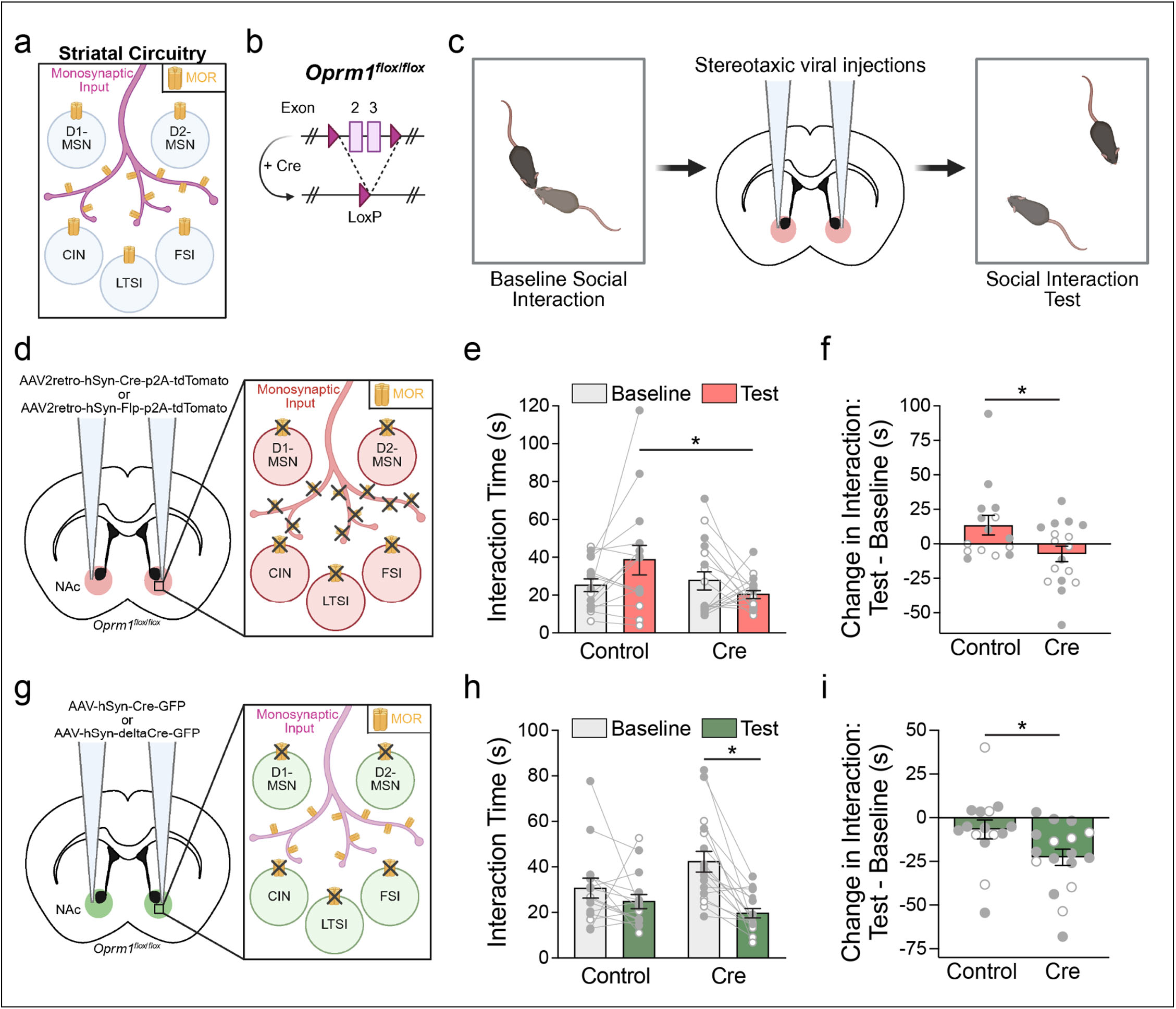
Reciprocal social interaction is impaired by conditional genetic knockout of Oprm1 from nucleus accumbens neurons. (a) Illustration of mu opioid receptor expression in striatal circuitry (b) Cre recombinase deletes exons 2 and 3 from the Oprm1^fl/fl^ allele. (c) Experimental timeline: baseline social interaction test followed by bilateral injection of AAV expressing Cre recombinase in the nucleus accumbens, with a social interaction test six weeks later. (d) Experimental strategy to delete Oprm1 from NAc neurons and presynaptic inputs using AAV2retro. (e) Total interaction time before and after expression of Cre recombinase with AAV2retro. (f) Change in social interaction expressed as a difference score between test and baseline sessions. (g) Experimental strategy to delete Oprm1 from all NAc neurons. (h) Total interaction time before and after expression of Cre expression with AAVdj. (i) Change in social interaction expressed as a difference score between test and baseline sessions. Data are mean +/- SEM (n=15-17); open and filled circles represent female and male mice, respectively. *p<0.05, ANOVA interaction followed by simple effect test (e and h) or main effect of group (f and i).

The vast majority (∼95%) of postsynaptic NAc neurons are medium spiny projection neurons, with different subpopulations expressing either the Drd1 or Drd2 dopamine receptor (D1-MSNs and D2-MSNs, respectively). We next crossed Oprm1^fl/fl^ mice with transgenic lines expressing Cre recombinase in D1-MSNs (Drd1-Cre) and/or D2-MSNs (Adora2a-Cre)^21^, which will remove Oprm1 expression from MSNs of each type across the striatum. To our surprise, social interaction was not significantly affected in mice carrying either Cre allele, or both Cre alleles, compared to Oprm1^fl/fl^ littermate controls lacking Cre (Fig. 2b-c). However, mice carrying both Cre alleles had a diminished locomotor response to acute morphine administration (Fig. 2d-e). Statistical analysis indicated a significant main effect of Genotype (F_3,31_ = 3.39, p = 0.030, η_p_^2^ = 0.25), demonstrating functional efficacy of this manipulation that resembles the effect of Oprm1 deletion from all forebrain GABA neurons^10^.

To corroborate this surprising effect of Oprm1 deletion throughout development from MSNs across the striatum, we conducted another experiment to locally delete Oprm1 from MSNs in the NAc of adult mice. We used a viral strategy to express Cre recombinase in MSNs using the CamKII promoter (Fig. 2f), because even though MSNs release GABA, they also express CaMKII at high levels because they are spiny projection neurons^22^. Control mice were similarly injected with AAV lacking Cre recombinase expression (AAV-CamKII-eGFP). As before, social interaction was assessed before and 6 weeks after stereotaxic viral injections (Fig. 2g). Importantly, social interaction following Oprm1 knock-out from NAc MSNs was not statistically different from baseline social interaction when comparing interaction time within subjects (Fig. 2h-i), corroborating that MORs on NAc MSNs are not required for typical social behavior.

**Figure 2.**
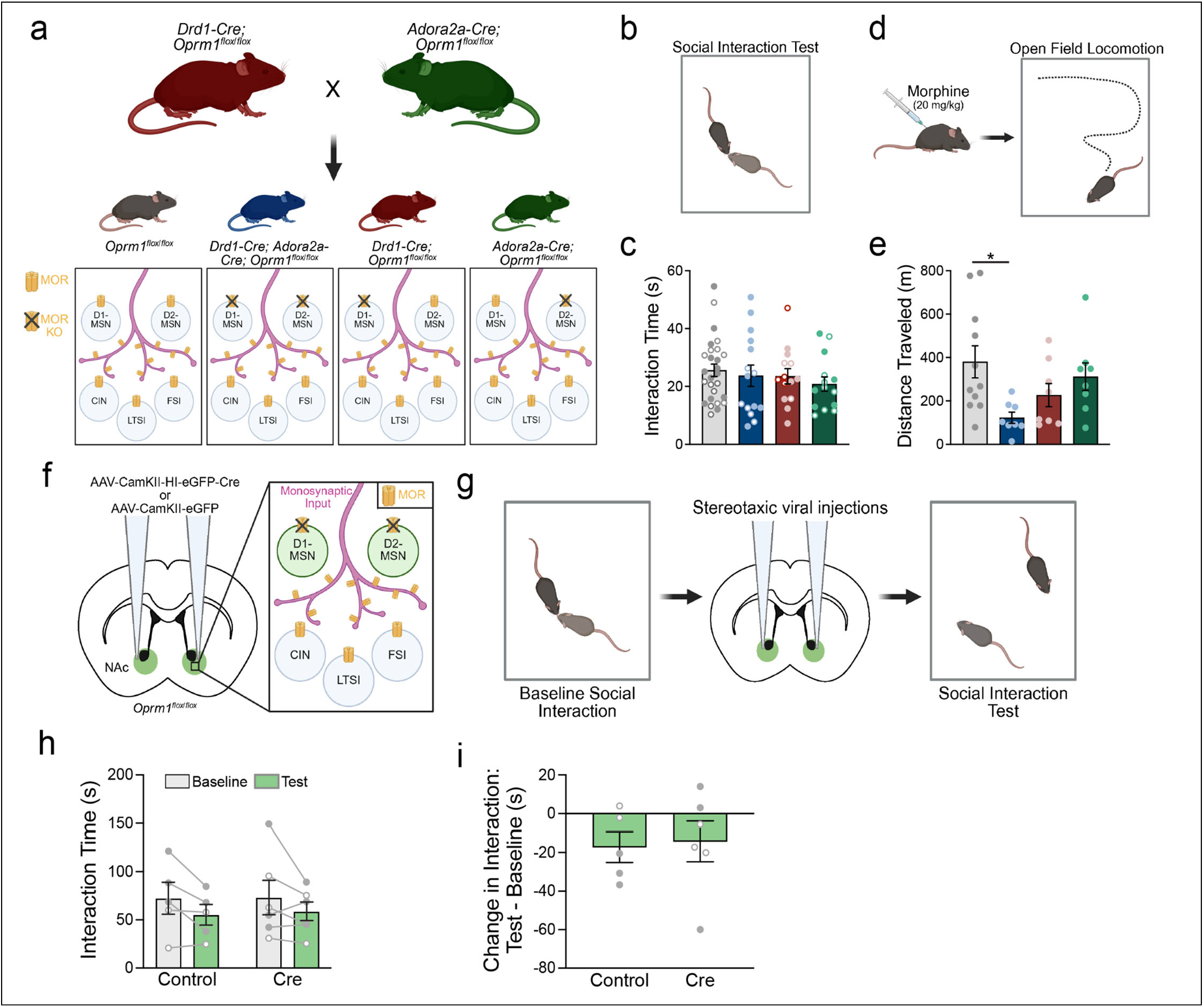
Conditional genetic knockout of Oprm1 from medium spiny projection neurons (MSNs) alters acute morphine sensitivity but not social interaction. (a) Breeding strategy used to generate Oprm1^fl/fl^ mice carrying Drd1-Cre and/or Adora2a-Cre (top), with a schematic illustration of genetic impact on MSNs (bottom). (b) Illustration of the social interaction test. (c) Social interaction testing of mice with Oprm1 deletion from both D1/D2-MSNs (blue), D1-MSNs (red), or D2-MSNs (green) compared to control Oprm1^fl/fl^ (gray) mice (n=14-27). (d) Illustration of locomotor test following acute morphine administration (e) Distance travelled after morphine administration in mice with Oprm1 deletion from both D1/D2-MSNs (blue), D1-MSNs (red), or D2-MSNs (green), compared to control Oprm1^fl/fl^ mice (gray), with significantly less distance travelled in mice carrying Drd1-Cre and Adora2a-Cre (n=8-11). (f) Experimental strategy to delete Oprm1 from NAc MSNs in adult mice, using an AAV to express Cre recombinase under control of the CamKII promoter. (g) Experimental timeline: baseline social interaction test followed by bilateral injection of AAV expressing Cre recombinase in the NAc followed by a social interaction test six weeks later. (h) Total interaction time before and after expression of Cre recombinase. (i) Change in social interaction expressed as a difference score between test and baseline session (n= 5-6 mice). Data are mean +/-SEM; open and filled circles represent female and male mice, respectively. *p<0.05, ANOVA followed by Fisher’s LSD post-hoc test (c and e).

Beyond MSNs, the NAc also contains multiple subtypes of local interneurons^18^. We first crossed Oprm1^fl/fl^ mice with a transgenic line expressing Cre recombinase in cholinergic interneurons, but this manipulation did not significantly affect social interaction (Extended Data Fig. 3). To target the remaining populations of inhibitory interneurons, which include LTSIs and FSIs (Fig. 3a), we used the Dlx enhancer sequence^23^ to control expression of Cre recombinase (AAV8-Dlx-iCre-tdTomato). We first used electrophysiology to evaluate the selectivity of this experimental approach for inhibitory interneurons. Cre recombinase can have catalytic activity when expressed in trace amounts, and very low levels of Cre expression are not reliably reported by fluorescent protein (i.e., tdTomato) expressed from the same viral vector^23^. We therefore co-expressed a second Cre-dependent reporter virus (AAV9-Ef1A-DIO-eYFP) to provide more sensitive readout of Cre enzymatic activity, and performed whole-cell current-clamp recordings from cells expressing eYFP to characterize their intrinsic membrane properties (Fig. 3b). The majority of these cells (∼75% of eYFP+ cells) exhibited electrophysiological characteristics of either FSIs^24^ or LTSIs^17^, two major inhibitory interneuron subtypes in the NAc^25^ (Fig. 3b). The remaining cells (∼25% of eYFP+ cells) exhibited MSN characteristics, consistent with prior reports that Cre expression with the Dlx enhancer sequence is highly preferential (but not perfectly selective) for inhibitory interneurons^26^.

Using fluorescent *in situ* hybridization (FISH), we also confirmed successful *Oprm1* deletion following Cre expression with the Dlx enhancer sequence. Towards this, we bilaterally injected either AAV8-Dlx-iCre-tdTomato or AAV8-Dlx-tdTomato (control virus) into the NAc of Oprm1^fl/fl^ mice (Fig. 3d). Six weeks later, we perfused brains, cryosectioned tissue, and performed FISH, probing for *Oprm1* RNA (Fig. 3e). Only 17% of tdTomato+ cells also expressed *Oprm1* transcripts in mice injected with AAV8-Dlx-iCre-tdTomato, compared to 64% of tdTomato+ cells in mice injected with AAV8-Dlx-tdTomato control virus (Fig. 3f; Extended Data Fig. 4). Having confirmed the efficacy and specificity of this manipulation, we proceeded with analysis of social behavior.

**Figure 3.**
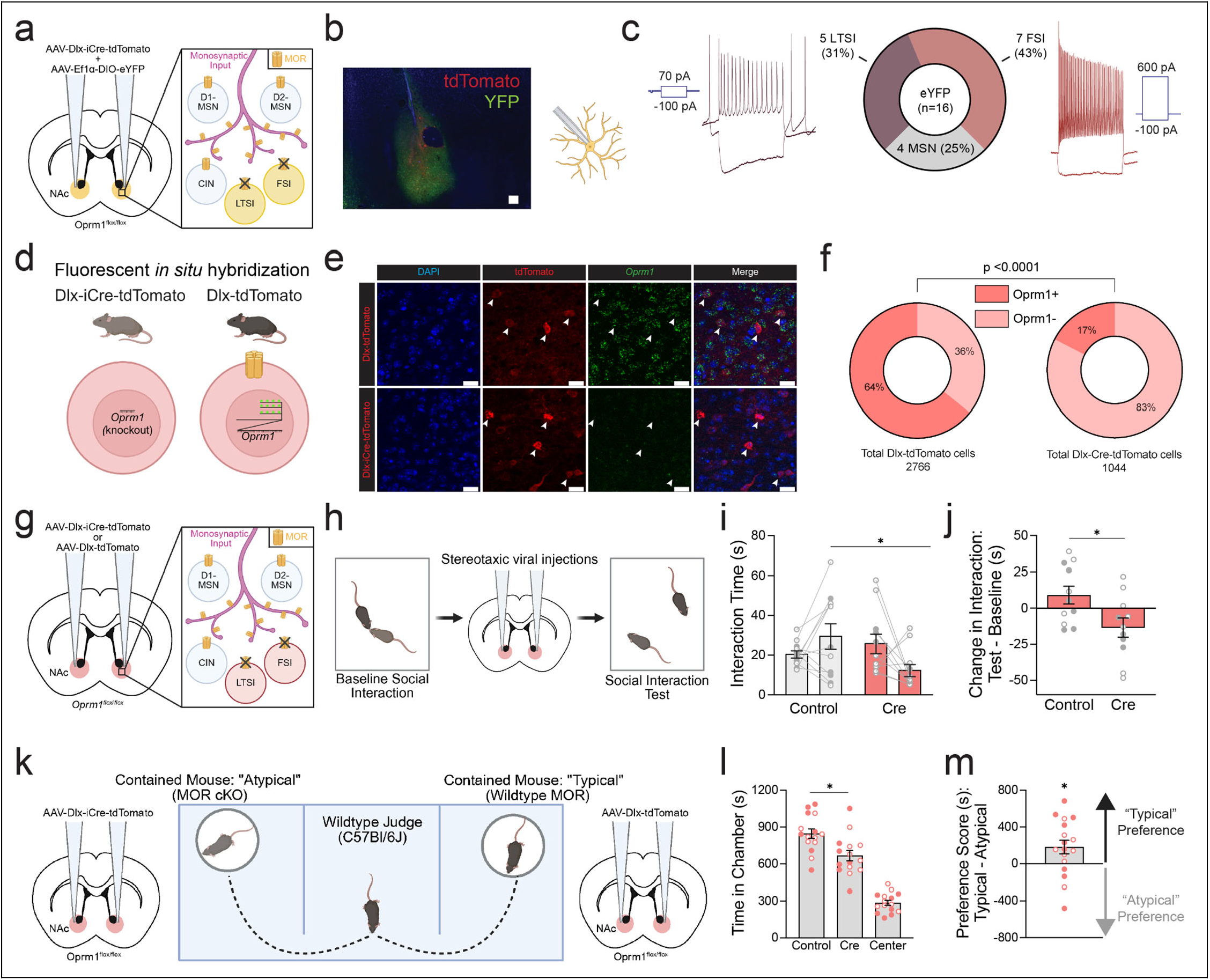
Social deficits following conditional genetic knockout of Oprm1 from nucleus accumbens inhibitory interneurons. (a) The Dlx enhancer sequence was used to drive viral expression of Cre recombinase, with Cre-dependent expression of eYFP to report enzymatic activity. (b) Representative image of virus expression in a coronal brain slice. (c) Whole-cell current-clamp recordings of cells expressing eYFP (n=16), which had electrophysiological characteristics of low threshold-spiking interneurons (LTSI), fast-spiking interneurons (FSI), or medium spiny projections neurons (MSN). (d) Fluorescent in situ hybridization was used to validate Oprm1 deletion following injection of Dlx-iCre-tdTomato, compared to Dlx-tdTomato control virus (e) Representative NAc images from mice injected with AAV-Dlx-tdTomato and AAV-Dlx-iCre-tdTomato. Scale bar=200um. (f) Proportion of tdTomato+ cells that are positive or negative for Oprm1 mRNA following injection with Dlx-tdTomato control virus (n=1044 cells from three mice) or Dlx-iCre-tdTomato virus (n=2766 cells from three mice). (g) Illustration of experimental strategy to knockout Oprm1 from LTSIs and FSIs concomitantly using the Dlx enhancer to drive Cre expression. (h) Timeline: baseline social interaction tested before bilateral NAc virus injection, followed by a social interaction test four weeks later. (i) Total interaction time before and after expression of Cre recombinase. (j) Change in social interaction expressed as a difference score between test and baseline session (n=11). (k) Illustration of real-time social preference test: C57Bl/6J judges (n=17) simultaneously engaged with confined social targets injected with Cre recombinase (left) or tdTomato (right). (l) Time spent in each chamber. (m) Preference score. Data are mean +/-SEM; open and filled circles represent female and male mice, respectively. *p<0.05, Fisher’s exact test (f), ANOVA interaction followed by simple effect test (i), ANOVA main effect of group (j), ANOVA main effect of chamber followed by Fisher’s LSD post-hoc test (l), or one-sample t-test (m).

After a baseline test of social interaction, we performed bilateral injections of AAV8-Dlx-iCre-tdTomato into the NAc of Oprm1^fl/fl^ mice (Fig. 3g-h and Extended Data Fig. 5). This manipulation significantly reduced social interaction when a test session was conducted after four weeks of viral expression, compared to an AAV8-Dlx-tdTomato control virus (Fig. 3i). Statistical analysis across baseline and test sessions indicated a significant Session x Virus interaction (F_1,18_ = 6.28, p = 0.022, η_p_^2^ = 0.26), which was evident when a difference score (test – baseline) was compared between genotypes (Fig. 3e). In a test of real-time social preference^27^ (Fig. 3k), C57BL/6J mice spent less time interacting with Oprm1^fl/fl^ mice that received AAV8-Dlx-iCre-tdTomato, compared to AAV8-Dlx-tdTomato control mice (Fig. 3l). Statistical analysis indicated a main effect of Chamber (F_2,42_ = 65.87, p < 0.0001, η_p_^2^ = 0.76), with a significant difference between time spent interacting with the “typical” (control) versus “atypical” (Cre; Fig. 3m). This mirrors previous characterization of Oprm1 constitutive knockout mice^16^, and provides further evidence for atypical social behavior after Oprm1 deletion from NAc inhibitory interneurons.

We also used distinct viral and genetic strategies to independently delete Oprm1 expression from either FSIs or LTSIs in the NAc. FSIs were targeted by injecting Oprm1^fl/fl^ mice with a virus using the S5E2 enhancer sequence^28^ to drive expression of Cre recombinase (Extended Data Fig. 6). LTSIs were targeted by injection of a Flp-dependent Cre virus into Oprm1^fl/fl^;Sst-ires-Flp mice (Extended Data Fig. 7). Neither of these manipulations had significant effects on social interaction, indicating that social behavior is only impaired by concomitant deletion of Oprm1 from all NAc inhibitory interneurons. This impairment did not generalize to fentanyl-conditioned place preference, which was intact following Oprm1 deletion from NAc inhibitory interneurons (Extended Data Fig. 8).

We next wanted to examine how Oprm1 deletion from inhibitory interneurons alters synaptic transmission onto the MSNs, as LTSIs and FSIs both provide potent inhibition onto MSNs, which are the principal output neurons of the NAc. Towards this, we bilaterally injected AAV-Dlx-iCre-tdTomato or AAV-Dlx-tdTomato into the NAc of Oprm1^fl/fl^;Drd2-eGFP mice (Fig. 4a). Six weeks later, we performed whole-cell voltage-clamp recordings from either D2-MSNs (GFP+) or putative D1-MSNs (GFP-). We first measured the amplitude of evoked inhibitory postsynaptic current (IPSC), which was normalized to the amplitude of evoked excitatory postsynaptic current (EPSC) recorded from the same cell, to calculate the ratio of inhibitory versus excitatory synaptic input (IPSC/EPSC ratio; Fig. 4b). Intriguingly, Oprm1 deletion from inhibitory interneurons reduced the IPSC/EPSC ratio in both D1- and D2-MSNs (Fig. 4c), indicating that removal of MORs from LTSIs and FSIs alters the overall balance of inhibitory and excitatory input onto both MSN subtypes in the NAc. However, there were no changes in paired-pulse ratio for either IPSCs or EPSCs (Fig. 4d-f), and no changes in the amplitude or frequency of spontaneous IPSCs (sIPSCs) measured between delivery of electrical stimuli (Fig. 4g-h). *Oprm1* deletion from NAc inhibitory interneurons thus appears to shift the balance between inhibitory and excitatory synaptic transmission, a change that may underlie the deficits in social behavior caused this manipulation.

**Figure 4.**
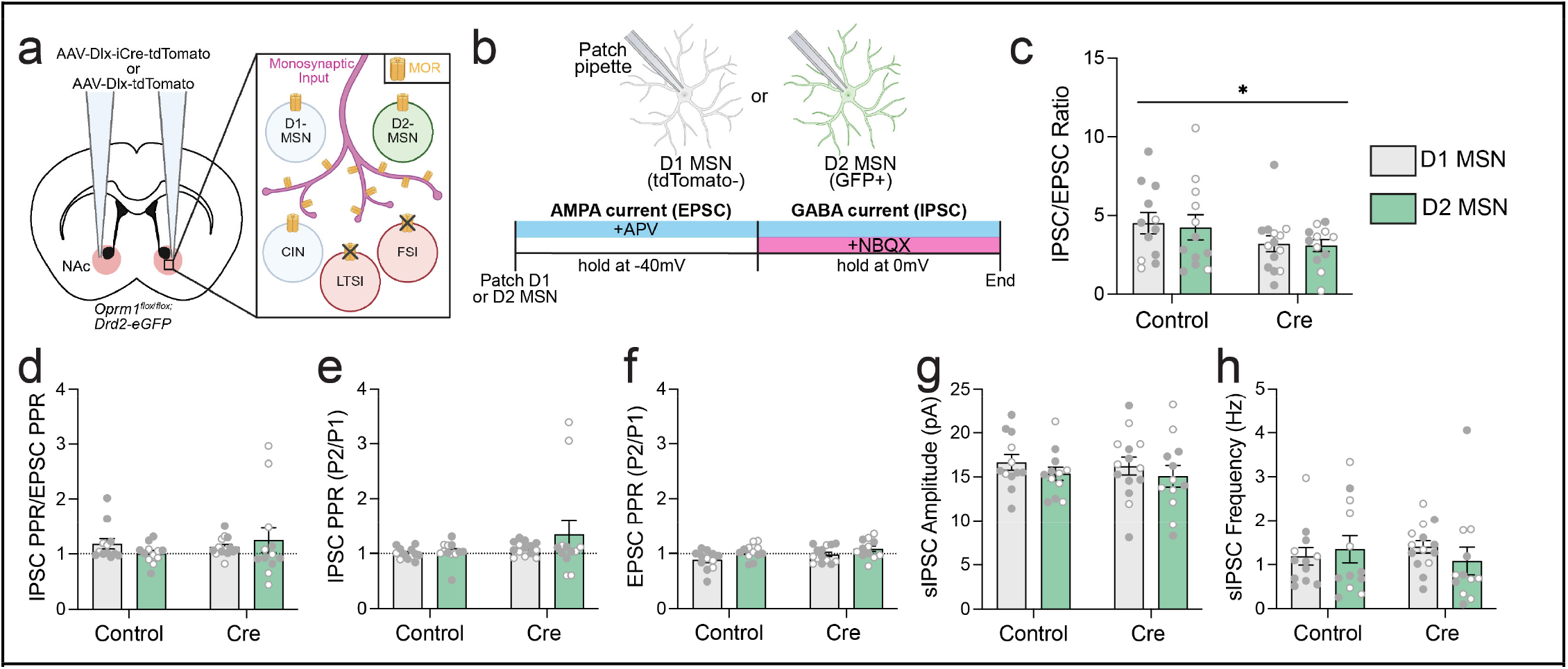
Oprm1 deletion from inhibitory interneurons alters the balanced between synaptic inhibition and excitation. (a) Experimental strategy using the Dlx enhancer sequence to drive viral expression of Cre recombinase Oprm1^fl/fl^;Drd2-eGFP mice. Control mice are injected with Dlx-tdTomato (no Cre expression). (b) Electrophysiological recording timeline of D2-MSNs (GFP+) or putative D1-MSNs (GFP-). First, AMPA current is isolated by holding the voltage at -40 mV while blocking NMDA receptors D-APV. Next, GABA current is isolated by increasing the holding voltage to 0 mV, and blocking AMPA receptors with addition of NBQX. Throughout recording, pairs of electrical stimuli were delivered (100 ms interval) to measure paired-pulse ratio (PPR), and spontaneous synaptic currents were recorded between stimuli. (c) The average IPSC/EPSC ratio in D1-and D2-MSNs in control and Cre mice. (d) IPSC PPR divided by EPSC PPR. (e) IPSC PPR. (f) EPSC PPR. (g) Spontaneous IPSC (sIPSC) amplitude. (h) Spontaneous IPSC frequency. Data are mean +/- SEM; open and filled circles represent individual cells from female and male mice, respectively. *p<0.05, ANOVA main effect of Group.

MORs are inhibitory G_i/o_-coupled receptors that cause neuronal hyperpolarization and reduced neurotransmitter release upon activation. Thus, one would hypothesize that removal of MORs would have an excitatory effect on cellular activity. To test this hypothesis, we used chemogenetics to directly activate NAc inhibitory interneurons while measuring social interaction, to determine if activation of these cells would phenocopy the behavioral effect of *Oprm1* deletion. Towards this, we stereotaxically injected a virus expressing the excitatory chemogenetic actuator, hM3Dq, driven by the Dlx enhancer sequence (AAV-Dlx-hM3Dq-tdTomato) to the NAc of wildtype mice (Fig. 5a). Using whole-cell current-clamp recordings from tdTomato+ cells, we verified this virus is preferentially expressed by inhibitory interneurons, with about 84% of tdTomato+ cells recorded having electrophysiological signatures of LTSIs or FSIs, while only 16% of tdTomato+ cells had MSN properties (Fig. 5b-c). We also performed cell-attached recordings to monitor spontaneous spiking activity, and verified that hM3Dq activation with deschloroclozapine (DCZ, 1 uM) significantly increased the firing rate of tdTomato+ cells (Fig. 5d-e).

**Figure 5.**
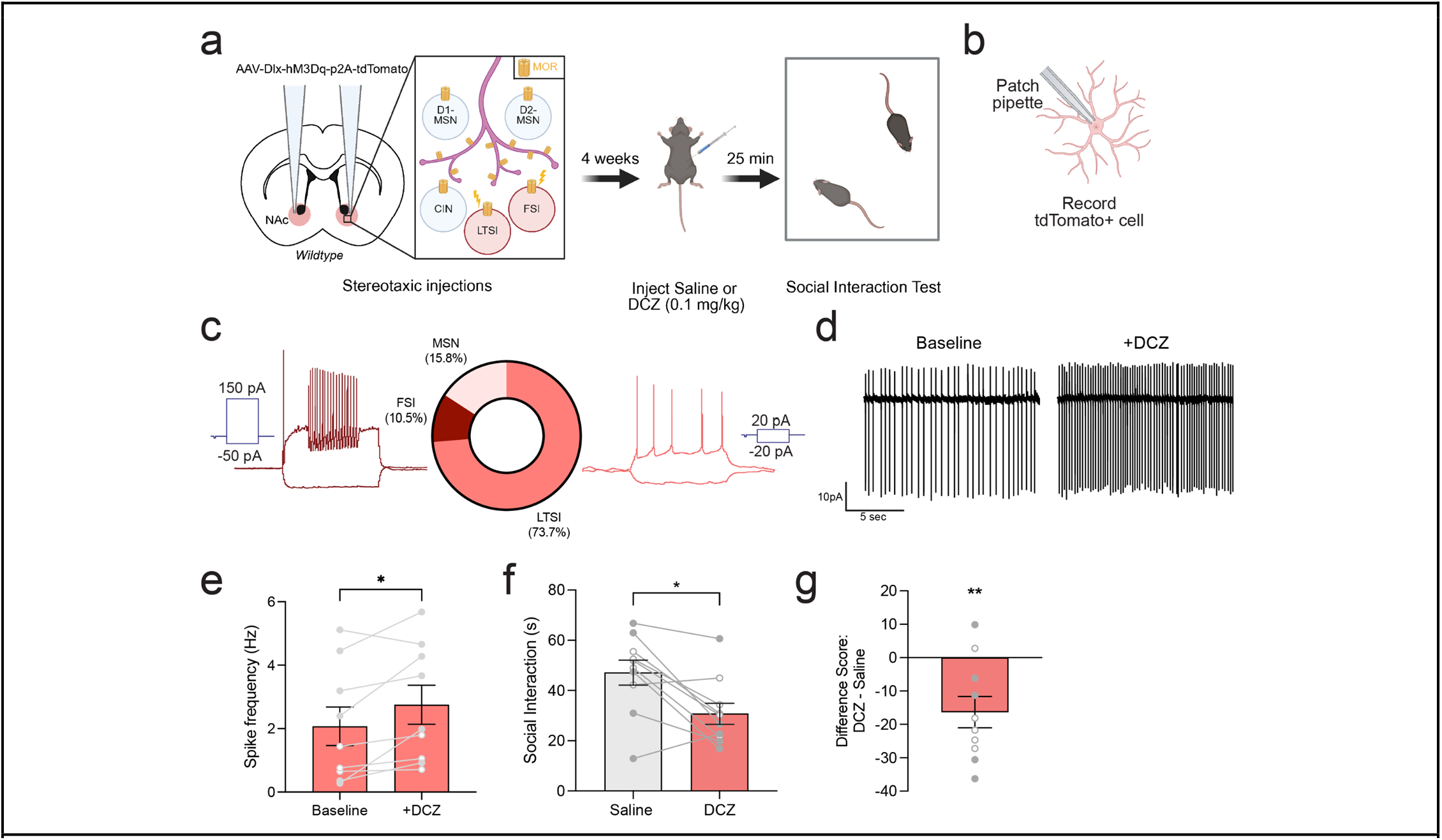
Chemogenetic activation of NAc inhibitory interneurons impairs reciprocal social interaction. (a) Experimental strategy using the Dlx enhancer sequence to drive viral expression of hM3Dq-tdTomato in wildtype mice. Social interaction testing occurs following treatment with either saline (control) or DCZ. (b) Electrophysiological recording of tdTomato+ cells for viral validation. (c) Whole-cell current-clamp recordings of cells expressing tdTomato (n=19), which had electrophysiological characteristics of low threshold-spiking interneurons (LTSI), fast-spiking interneurons (FSI), or medium spiny projections neurons (MSN). (d) Sample trace of cell-attached recording from tdTomato+ cell at baseline and after washing on DCZ (1 uM). (e) Average spike frequency (Hz) from cell-attached recordings of tdTomato+ cells at baseline and after washing on DCZ. (f) Total interaction time of Dlx-hM3Dq mice following treatment with saline or DCZ (g) Change in social interaction expressed as a difference score between test and baseline session (n=10). Data are mean +/- SEM; open and filled circles represent female and male mice, respectively. *p<0.05, paired t-test (e-f) or one sample t test (g).

To evaluate the behavioral effects of this chemogenetic manipulation, mice received bilateral injection of AAV-Dlx-hM3Dq-tdTomato in to the NAc. Four weeks later, we conducted two social interaction tests following injection of either saline or DCZ (0.1 mg/kg), with one week separating test sessions and treatment order counterbalanced between animals. Compared to saline, treatment with DCZ significantly decreased social interaction time compared to saline treatment (Fig. 5f-g). This indicates that chemogenetic activation of inhibitory interneurons in the NAC is sufficient to produce social behavior deficits, mimicking the effect of Oprm1 deletion from this same population of cells.

## DISCUSSION

Our results pinpoint NAc inhibitory interneurons as a critical cellular locus where mu opioid receptor expression supports typical social behavior. Genetic deletion of *Oprm1* from all inhibitory interneurons, including both LTSIs and FSIs, phenocopied the social behavioral deficits observed with pan-neuronal knockout of *Oprm1*, indicating that MOR signaling on inhibitory interneurons is critical for typical social behavior. Mice with Oprm1 deletion from inhibitory interneurons exhibited less social interaction themselves, and also appeared to be perceived differently by conspecifics, as wildtype mice were less inclined to interact with “atypical” Oprm1 conditional knockout mice compared to “typical” control mice. We previously observed these social behavior deficits in constitutive Oprm1 knockout mice^16^, and the present results show that conditional Oprm1 deletion from NAc inhibitory interneurons is sufficient to generate these social phenotypes.

Both FSIs and LTSIs provide inhibitory synaptic input to MSNs^17, 24, 25^, thereby controlling output from NAc projection neurons. For normal expression of social behavior, mu opioid receptor activation may be necessary to decrease the activity of inhibitory interneurons, and release MSNs from synaptic inhibition. When mu opioid receptors are deleted from only FSIs or LTSIs, intact expression by the other inhibitory interneuron subtype appears to be sufficient for normal expression of social behavior. Simultaneous deletion of the mu opioid receptor from both FSIs and LTSIs may tip the system beyond a critical threshold that is not reached when either cell type is manipulated in isolation, likely reflecting a difference between partial versus total lack of synaptic disinhibition of MSN output during social interaction. While we observed Cre activity in a small number of MSNs using the Dlx enhancer sequence, we believe MSN expression is unlikely to contribute to social behavior deficits, because there was no effect on social behavior following conditional Oprm1 deletion from MSNs using genetic or viral strategies.

We hypothesized that MOR deletion from inhibitory interneurons would increase their activity, resulting in greater inhibition onto downstream MSNs and ultimately impairing social behavior. Indeed, we found that direct chemogenetic activation of NAc interneurons using hM3Dq (an excitatory DREADD) was sufficient to reduce social interaction, in a manner similar to MOR deletion from inhibitory interneurons. However, when we performed electrophysiological recordings to measure inhibitory and excitatory synaptic transmission onto D1- and D2-MSNs after MOR deletion from inhibitory interneurons, we observed a decrease in the inhibition:excitation ratio. In theory, this change could represent either a decrease in inhibitory input onto NAc MSNs, or an increase in excitatory input onto NAc MSNs, but we did not find any evidence of a change in the probability of presynaptic release for either glutamate or GABA onto MSNs. We also did not observe any change in sIPSC amplitude, a measure sensitive to postsynaptic changes to inhibitory neurotransmission. However, there are several possible explanations for this surprising effect on NAc circuitry. Foremost, LTSIs and FSIs co-release other neurotransmitters in addition to GABA, including neuropeptide Y (NPY), somatostatin, and nitric oxide^29^. Of note, NPY release by LTSIs in the NAc has been shown to promote social behavior^30^, and can have either excitatory or inhibitory postsynaptic effects depending on the NPY receptor subtype engaged. Thus, perhaps MOR deletion from inhibitory interneurons is altering neuropeptide release to change the overall balance between synaptic inhibition and excitation onto NAc MSNs in a manner independent from direct release of GABA. We also note a limitation in our analysis of synaptic transmission, which was the use of conventional electrical stimulation to activate all synaptic inputs regardless of source. For inhibitory transmission, this means we monitored the response to all sources of inhibitory synaptic input, which include collateral connections between MSNs as well as input from LTSIs and FSIs^31^. Future studies could use paired whole-cell patch-clamp recordings or optogenetic strategies to isolate synaptic input from specific sources, and quantify the impact of Oprm1 deletion more precisely.

One fascinating aspect of our results is that the same mu opioid receptor manipulations that disrupted social behavior had no apparent impact on sensitivity to exogenous opioid administration. Conversely, acute morphine sensitivity was diminished by deletion of the mu opioid receptor from all MSNs, but this manipulation had no effect on social behavior. This dissociation is highly relevant to the therapeutic potential of mu opioid receptor stimulation to enhance sociability^32^. It points toward the possibility that engaging mu opioid receptors in a circuit-specific fashion could circumvent the risks of misuse and abuse associated with conventional mu opioid receptor agonists, which have indiscriminate action across cell types and brain circuits. Our findings thus have important implications for both the pathophysiology and treatment of neuropsychiatric conditions that involve deficits in affiliative social behavior.

## Acknowledgments

Research reported in this publication was supported by the University of Minnesota’s MnDRIVE (Minnesota’s Discovery, Research and Innovation Economy) initiative (E.M.L. & P.E.R.), as well as grants from the National Institutes of Health: DA007234 (C.T.), MH122094 (C.T.), DA060664 (J.L.S.), DA052624 (E.M.L.), MH119569 (E.M.F.V. & P.E.R.), DA048946 (P.E.R.), and DA065635 (P.E.R.). Some viral vectors used in this study were generated by the University of Minnesota Viral Vector and Cloning Core and the Center for Neural Circuits in Addiction (P30 DA048742). We thank the University of Minnesota Mouse Behavior Core for use of their facilities to conduct behavioral tests.

## METHODS AND MATERIALS

### Subjects

Experiments were performed with female and male C57Bl/6J mice (The Jackson Laboratory strain #000664); Oprm1 conditional knockout mice (The Jackson Laboratory strain #030074)^19^; Drd1-Cre BAC transgenic FK15022 (Mutant Mouse Resource & Research Centers #029178-UCD)^33^; Adora2a-Cre BAC transgenic KG13923 (Mutant Mouse Resource & Research Centers #036158-UCD)^34^; ChAT-ires-Cre (The Jackson Laboratory strain #018957)^35^; Sst-ires-FLP (The Jackson Laboratory strain #031629)^36^; and Drd2-eGFP (Mutant Mouse Resource & Research Centers #036931-UCD)^37^. All genetically modified strains were maintained on a C57Bl/6J genetic background. Mice were housed in groups of 2-5 per cage, on a 12-hour light cycle (0600h–1800h) at ∼23° C with ad libitum access to food and water. Baseline tests of social interaction were conducted at ∼6 weeks of age, followed by stereotaxic surgery at ∼7 weeks of age. Experimental procedures were conducted between 1000h–1600h and were approved by to the University of Minnesota Institutional Animal Care and Use Committee and observed the NIH Guidelines for the Care and Use of Laboratory Animals.

### Adeno-Associated Viral Vectors and Stereotaxic Surgery

AAV2retro-hSyn-Cre-P2A-tdTomato and AAV2retro-hSyn-Flp-P2A-tdTomato were generated by the University of Minnesota Viral Vector and Cloning Core; rAAV2-retro helper (Addgene plasmid # 81070) was a gift from Alla Karpova & David Schaffer^20^. AAVdj-hSyn-Cre-GFP and AAVdj-hSyn-ΔCre-GFP were obtained from the Stanford University Gene Vector and Virus Core, as previously described^21^. AAV9-CamKII-H1-GFP-Cre-WPRE-SV40 was a gift from James M. Wilson (Addgene viral prep # 105551-AAV9). AAV9-CamKII-GFP was a gift from Bryan Roth (Addgene viral prep # 50469-AAV9). AAV8-mDlx-iCre-tdTomato and AAV8-mDlx-tdTomato were generated by the University of Minnesota Viral Vector and Cloning Core, using a backbone (Addgene plasmid # 83900) that was a gift from Gordon Fishell^23^. AAV9-Ef1a-DIO-eYFP was a gift from Karl Deisseroth (Addgene viral prep # 27056-AAV9). AAV9-S5E2-tdtomato and AAV9-S5E2-Cre-P2A-tdTomato were generated by the University of Minnesota Viral Vector and Cloning Core, using a backbone (Addgene plasmid # 135630) that was a gift from Jordane Dimidschstein^28^. AAVdj-CAG-fDIO-GFP-Cre was generated by the University of Minnesota Viral Vector and Cloning Core. AAV9-hDlx-GqDREADD-dTomato-Fischell-4 was a gift from Gordon Fishell^23^ (Addgene viral prep # 83897-AAV9).

For intracranial virus injection, mice were anesthetized with a ketamine:xylazine cocktail (100:10 mg/kg). A small hole was drilled above target coordinates for the nucleus accumbens (AP +1.35, ML +1.10; DV -4.40), targeting the border between the core and shell subregions, thus infecting cells in both subregions. A small volume of virus (0.3-0.5 ul) in a glass needle was infused at 0.1 ul/min, and the injection needle was left in place for at least 5 min after the end of the infusion before being withdrawn. After surgery, mice were given 500 uL saline and 5 mg/kg carprofen (subcutaneous) daily for 3 days. Mice recovered for at least four weeks after intracranial injection before testing the behavioral consequences of viral manipulations, to provide time for both genetic recombination and turnover of pre-existing mu opioid receptor protein. For experiments involving the AAV2retro serotype, mice recovered for six weeks after virus injection before the social interaction test, providing additional time for retrograde transport to sources of monosynaptic input to the NAc.

### Social Behavior Assays

Tests of reciprocal social interaction and real-time social preference were conducted as previously described^16^. Animals were moved to the behavior room one hour before testing. All experiments were conducted at 60-70 luminosity, and at a temperature equal to that of the animal housing facility. Experimental sessions were video recorded and hand-scored by researchers blind to experimental conditions. All tests of social behavior involved novel social partners that were not siblings or cage mates.

For reciprocal social interaction, the test apparatus was an opaque white rectangular box with 1 cm of fresh corn cob bedding on the floor. Experimental mice were introduced to an age- and sex-matched stimulus mouse in the testing apparatus for 10 min. Each stimulus mouse was a novel C57Bl/6J mouse, and was only used as a stimulus mouse for a single test session. Video recordings of various social behaviors exhibited by experimental and stimulus mice were hand scored by a blinded experimenter using Button Box 5.0 (Behavioral Research Solutions). Social interaction time was quantified as the sum of nose-nose interaction (direct investigation of orofacial region), huddling (stationary sitting next to partner), social exploration (anogenital investigation, social sniffing outside of orofacial region, social grooming), and following^38^. No aggressive interactions were documented in any group.

The real-time social preference test is based on a published protocol^27^ that allows a wild-type C57Bl/6J “judge” to choose between interacting with a “typical” and an “atypical” mouse. The test apparatus was a white plastic rectangular box (25” x 15”x 8”) consisting of three interconnected chambers. Prior to testing, judges were habituated to the empty apparatus for 10 minutes of free exploration. After habituation, two wire cups were placed in either end chamber: one wire cup contained an Oprm1^fl/fl^ mouse with bilateral injection of AAV8-mDlx-tdTomato in the nucleus accumbens (“typical”), and the other wire cup contained an Oprm1^fl/fl^ mouse with bilateral injection of AAV8-mDlx-iCre-tdTomato in the nucleus accumbens (“atypical”). Judges were then allowed to freely explore the chamber for 30 minutes. Test sessions were recorded by a video camera and the time the target mouse spent in each chamber was measured by ANY-maze tracking software. After each test, the entire apparatus was cleaned with 70% ethanol.

### Behavioral Responses to Exogenous Opioid Administration

Measurement of open field locomotion after morphine administration was performed as previously described^16^. We tested open-field locomotor activity in a clear plexiglass arena (ENV-510, Med Associates) housed within a sound-attenuating chamber. The location of the mouse within the arena was tracked in two dimensions by arrays of infrared beams, connected to a computer running Activity Monitor software (Med Associates). Mice were habituated to the chamber for one hour the day before initiating drug treatment. The next day, animals were tested in the open field chamber after injection of saline (s.c.), followed by morphine (2.0-20 mg/kg) on subsequent days. The session duration varied as a function of dose: 60 mins (saline and 2 mg/kg), 90 mins (6.32 mg/kg), or 120 mins (20 mg/kg).

Place conditioning with fentanyl was performed as previously described^39^, using activity test chambers (Med Associates, 11 x 11 x 11 in.) with two types of removable floor tiles: transparent plastic with a rough texture and black plastic with a smooth texture. A five-day protocol was used which included a 20 min baseline session on day 1 that included both contexts; three conditioning days consisting of one 30 min saline (5 mL/kg s.c.) conditioning session in the morning paired to one context and one 30 min fentanyl (0.04 mg/kg s.c.) conditioning session paired to the other context in the afternoon; and a 20 min test session on day 5 containing both contexts. Baseline preference was used to assign fentanyl administration to either a smooth black or rough transparent floor and the opposite floor for saline, so that the group average baseline preference was approximately 50%. Mice with a baseline preference greater than 75% for either floor were excluded. Conditioning sessions were at least three hours apart. Injections of saline or fentanyl were given 5 min prior to placement in activity chambers on conditioning days.

### Brain Slice Electrophysiology

Coronal brain slices (240 µm) containing the nucleus accumbens were prepared as previously described^16^. Mice were anesthetized with isoflurane and decapitated, brains quickly removed and placed in ice-cold cutting solution containing (in mM): 228 sucrose, 26 NaHCO3, 11 glucose, 2.5 KCl, 1 NaH2PO4-H2O, 7 MgSO4-7H20, 0.5 CaCl2-2H2O. Slices were cut by adhering the brain surface to the stage of a vibratome (Leica VT1000S), and allowed to recover in a submerged holding chamber with artificial cerebrospinal fluid (aCSF) containing (in mM): 119 NaCl, 26.2 NaHCO3, 2.5 KCl, 1 NaH2PO4-H2O, 11 glucose, 1.3 MgSO4-7H2O, 2.5 CaCl2-2H2O. Slices recovered in warm aCSF (33°C) for 15 min and then equilibrated to room temperature (∼25°C) for at least one hour before use. Slices were transferred to a submerged recording chamber and continuously perfused with aCSF at a rate of 2 mL/min at room temperature. All solutions were continuously oxygenated (95% O2/5% CO2).

Whole-cell recordings from fluorescent neurons in the nucleus accumbens were obtained under visual control using an Olympus BX51W1 microscope under IR-DIC optics. Current-clamp recordings were made with borosilicate glass electrodes (3-5 MΩ) filled with (in mM): 120 K-Gluconate, 20 KCl, 10 HEPES, 0.2 EGTA, 2 MgCl2, 4 ATP-Mg, 0.3 GTP-Na (pH 7.2-7.3). For voltage-clamp recording, electrodes were filled with (in mM): 120 CsMeSO_4_, 15 CsCl, 10 TEA-Cl, 8 NaCl, 10 HEPES, 5 QX-314, 4 ATP-Mg, 1 EGTA, 0.3 GTP-Na (pH 7.2-7.3). For cell-attached recordings, electrodes were filled with aCSF. Recordings were performed using a MultiClamp 700B amplifier (Molecular Devices), filtered at 2 kHz, and digitized at 10 kHz. Data acquisition and analysis were performed online using Axograph software. Series resistance was monitored continuously, and experiments were discarded if resistance changed by >20%. All reported holding potentials are corrected for a liquid junction potential of ∼10 mV.

Current-clamp recordings were used to characterize the intrinsic membrane properties of cells expressing different viral constructs. Fluorescent cells were injected with a series of current steps (1 sec duration) ranging from -100 to +600 pA. Maximum firing rate was calculated as the average maximum firing rate over the 1 sec step that could be sustained without inducing a depolarization block. Action potential half-width was measured at the half-way point between threshold and peak.

To measure evoked synaptic transmission during voltage-clamp recording, electrical stimulation was delivered using a bipolar electrode (50 us pulse width), with paired pulses (100 ms interval) delivered every 30 s. The ratio between synaptic inhibition and excitation (I/E ratio) was measured as previously described^21^. First, AMPA receptor-mediated transmission was isolated by holding the membrane potential at -40 mV, the reversal potential for GABA_A_ receptor-mediated current under our experimental conditions. while blocking NMDA receptors with D-APV (50 uM). Cells were subsequently depolarized to 0 mV and NBQX (10 uM) was applied, blocking AMPA receptors and leaving a residual current mediated by monosynaptic inhibition through GABA_A_ receptors. D-APV (50 uM) was present throughout the experiment to block NMDA receptors. The inhibition/excitation ratio was calculated by dividing peak GABA_A_ receptor current by peak AMPA receptor current for the first of the paired pulses, and sIPSCs were recorded at 0 mV during the period between electrical stimuli. Cell-attached recordings were used to validate the physiological response to activation of virally-expressed chemogenetic actuator hM3Dq. The spontaneous spiking of fluorescent cells was recorded in the presence of glutamate receptor antagonists (50 uM D-APV and 10 uM NBQX), GABA_A_ receptor antagonist (50 uM picrotoxin), and acetylcholine receptor antagonists (20 uM mecamylamine and 10 uM atropine), as previously described^17^. After recording baseline spiking, DCZ (1 uM) was washed on for 15 minutes.

### Immunohistochemistry and Epifluorescent Microscopy

Mice were deeply anesthetized using sodium pentobarbital (Fatal-Plus, Vortech Pharmaceuticals) and transcardially perfused with ice cold 0.01 M PBS followed by ice cold 4% PFA in 0.01 M PBS. Brains were removed and post-fixed 24 hours in 4% PFA in PBS. The following day, brains were rinsed briefly with 0.01 M PBS and sectioned in the coronal plane at 50 um. Tissue sections were blocked for 1 hour in blocking buffer (2% NHS, 0.2% triton x 100, and 0.05% Tween20 in 0.01 M PBS) and exposed to rabbit anti-GFP (1:1000, Abcam #ab290) or mouse anti-mCherry (1:1000, Abcam #125096), diluted in blocking buffer. After 24 hours at 4° C, sections were rinsed in wash buffer (Tris-buffered saline with 0.1% Tween20) and exposed to anti-Rabbit A488 (1:1000, Abcam #ab150073) or anti-Mouse A647 secondary antibodies (1:1000, Abcam #ab150115) overnight at 4° C. Stained tissue sections were imaged on a Keyence Microscope and images were compiled into a reference atlas.

### Fluorescent *in situ* Hybridization and Confocal Microscopy

Mice were deeply anesthetized with isoflurane and transcardially perfused with 10mL of ice cold 0.01 M PBS followed by 10 mL of ice cold 4% PFA in 0.01 M PBS. Brains were dissected and fixed overnight in 4% PFA in PBS, and then transferred to 10% sucrose in PBS. Twenty-four hours later brains were flash frozen in 2-methylbutane (∼ -25°C) and stored in a -80°C until cryosectioning. For cryosectioning, 16 um coronal slices containing the NAc were collected and adhered to pre-chilled slides (ColorFrost Plus). Following sectioning, fluorescent in situ hybridization was performed using the RNAscope v2 kit. The Oprm1-O4 probe (ACD Bio #544731) was used since this probe encompasses exons 2 and 3 of Oprm1, which are flnaked by LoxP sites in our Oprm1^fl/fl^ mouse line. The protease-free protocol using Pretreat Pro reagent for fixed/frozen tissue was followed explicitly, and 4% PFA was used for post-fixation of slides. Opal 690 (1:1000) was used to fluorescently label Oprm1, and endogenous tdTomato fluorescence was preserved throughout the RNAscope protocol. Following the RNAscope protocol, confocal imaging was performed using a Stellaris 8 DIVE confocal microscope (Leica). *Oprm1* RNA and tdTomato colocalization were quantified using HALO computer software (v3.6.4134.396).

### Statistical Analyses

Similar numbers of male and female animals were used in all experiments, with samples size indicated in figure legends and illustrated by individual data points in figures, with a visual distinction between data points from females (open circles) and males (filled circles). Sex was included as a variable in factorial ANOVA models analyzed using IBM SPSS Statistics v24, with repeated measures on within-subject factors. Main effects of sex and interactions involving sex were not significant unless noted otherwise. For main effects or interactions involving repeated measures, the Huynh-Feldt correction was applied to control for potential violations of the sphericity assumption. This correction reduces the degrees of freedom, resulting in non-integer values. Significant ANOVA interactions were decomposed by analyzing simple effects (i.e., the effect of one variable at each level of the other variable), and visualized by comparing difference scores between groups^40^. Significan main effects were analyzed using Fisher’s LSD post-hoc tests. Effect sizes are expressed as partial eta-squared (η_2_^2^) values. The Type I error rate was set to α = 0.05 (two-tailed) for all comparisons. All summary data are displayed as mean + SEM.

---

**Extended Data Figure 1.**
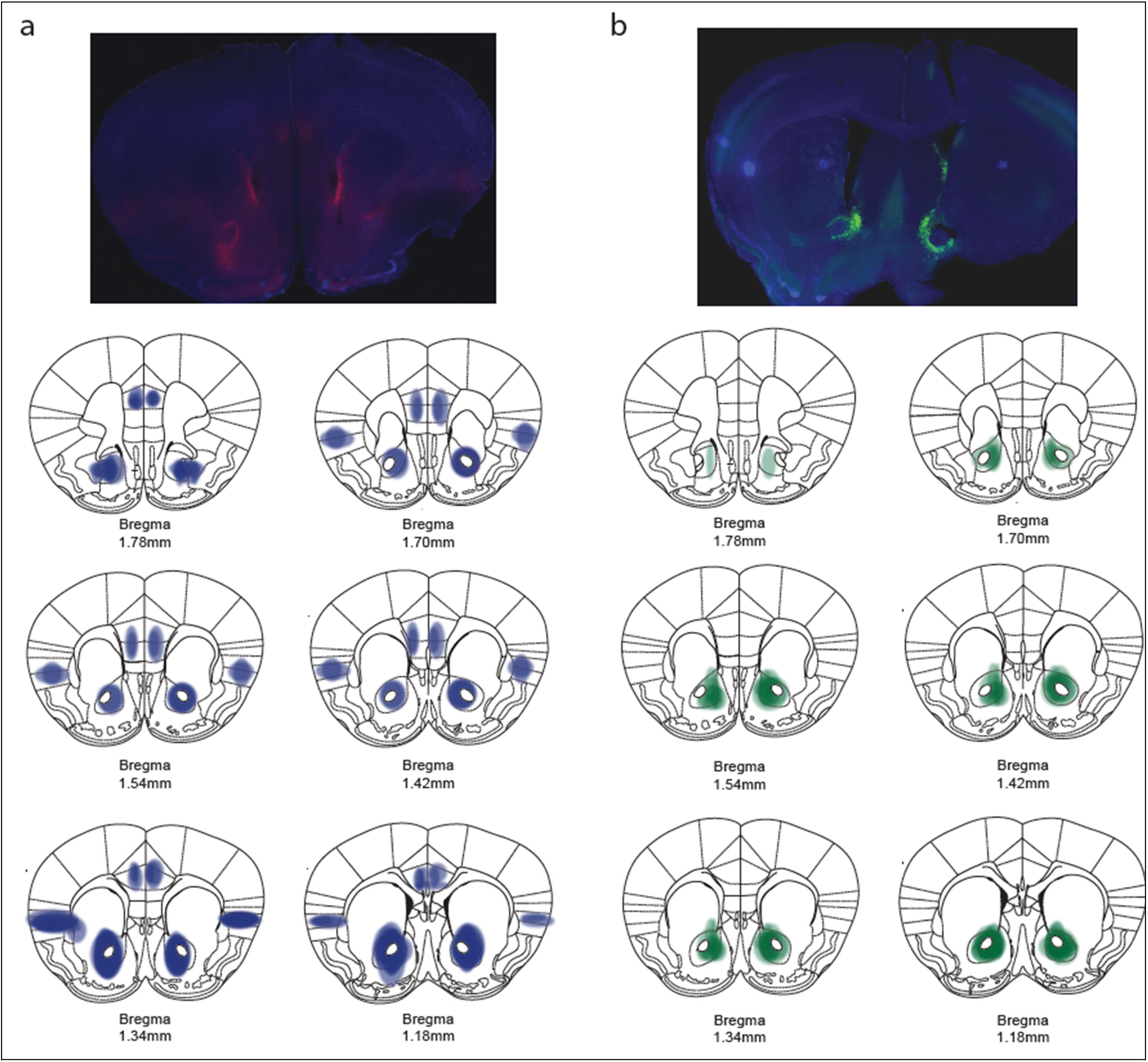
Viral expression patterns following nucleus accumbens injection of AAV2retro-hSyn-Cre-P2A-tdTomato (a) or AAVdj-hSyn-Cre-GFP (b). Representative images (top) and expression patterns in different coronal planes (bottom); retrograde transport to the medial prefrontal cortex and insular cortex was evident with the AAV2retro serotype (a).

**Extended Data Figure 2.**
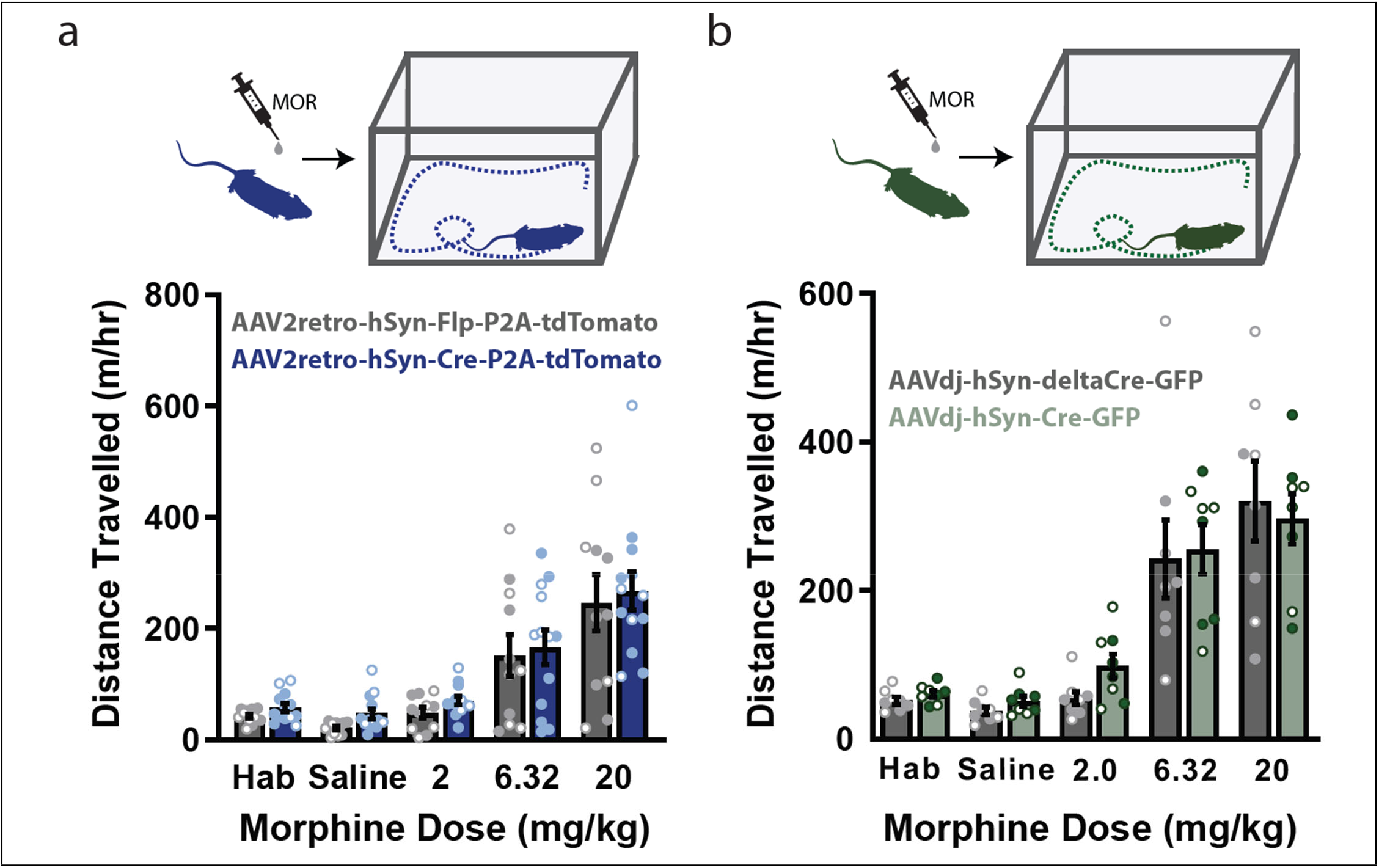
Conditional genetic knockout of Oprm1 from nucleus accumbens neurons does not affect behavioral sensitivity to acute morphine exposure. Distance travelled in the open field following injection of different morphine doses is shown for Oprm1fl/fl mice after expression of Cre recombinase with AAV2retro (a) or AAVdj (b), compared to a control virus. Data are mean +/-SEM (n = 8-13); open and filled circles represent female and male mice, respectively.

**Extended Data Figure 3.**
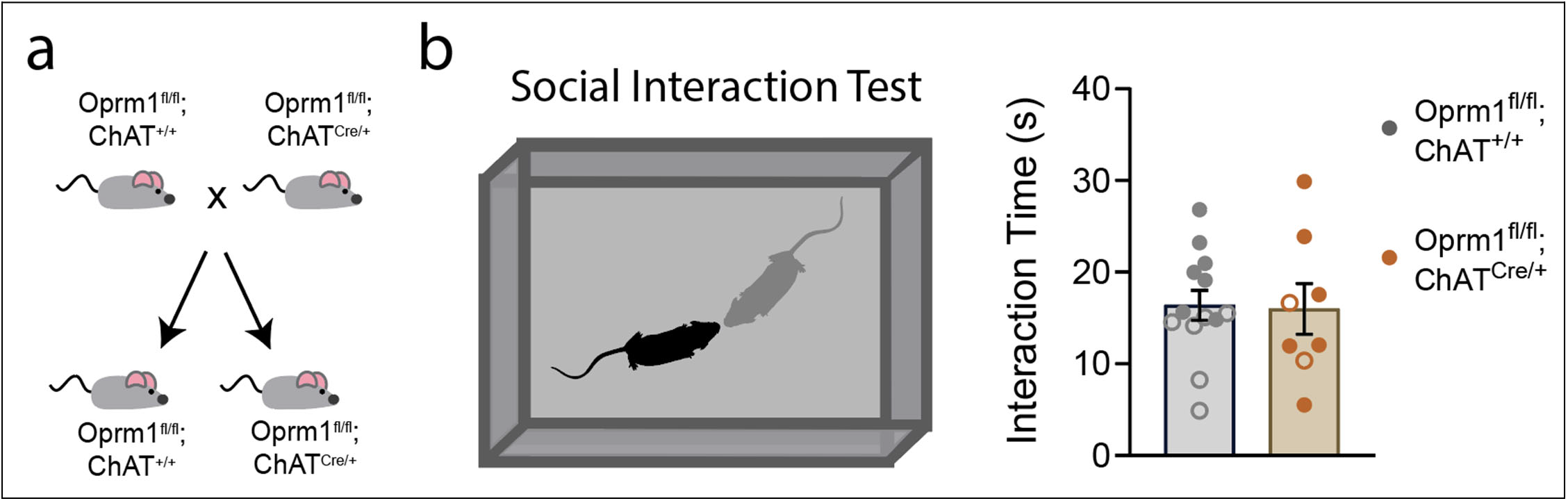
Conditional genetic knockout of Oprm1 from cholinergic neurons does not affect social interaction. (a) Breeding strategy used to generate Oprm1fl/fl mice expressing Cre recombinase from the choline acetyltransferase (ChAT) locus. (b) Illustration of the social interaction test (left), with no change in social interaction time between genotypes (right; n=8-13). Data are mean +/-SEM; open and filled circles represent female and male mice, respectively.

**Extended Data Figure 4.**
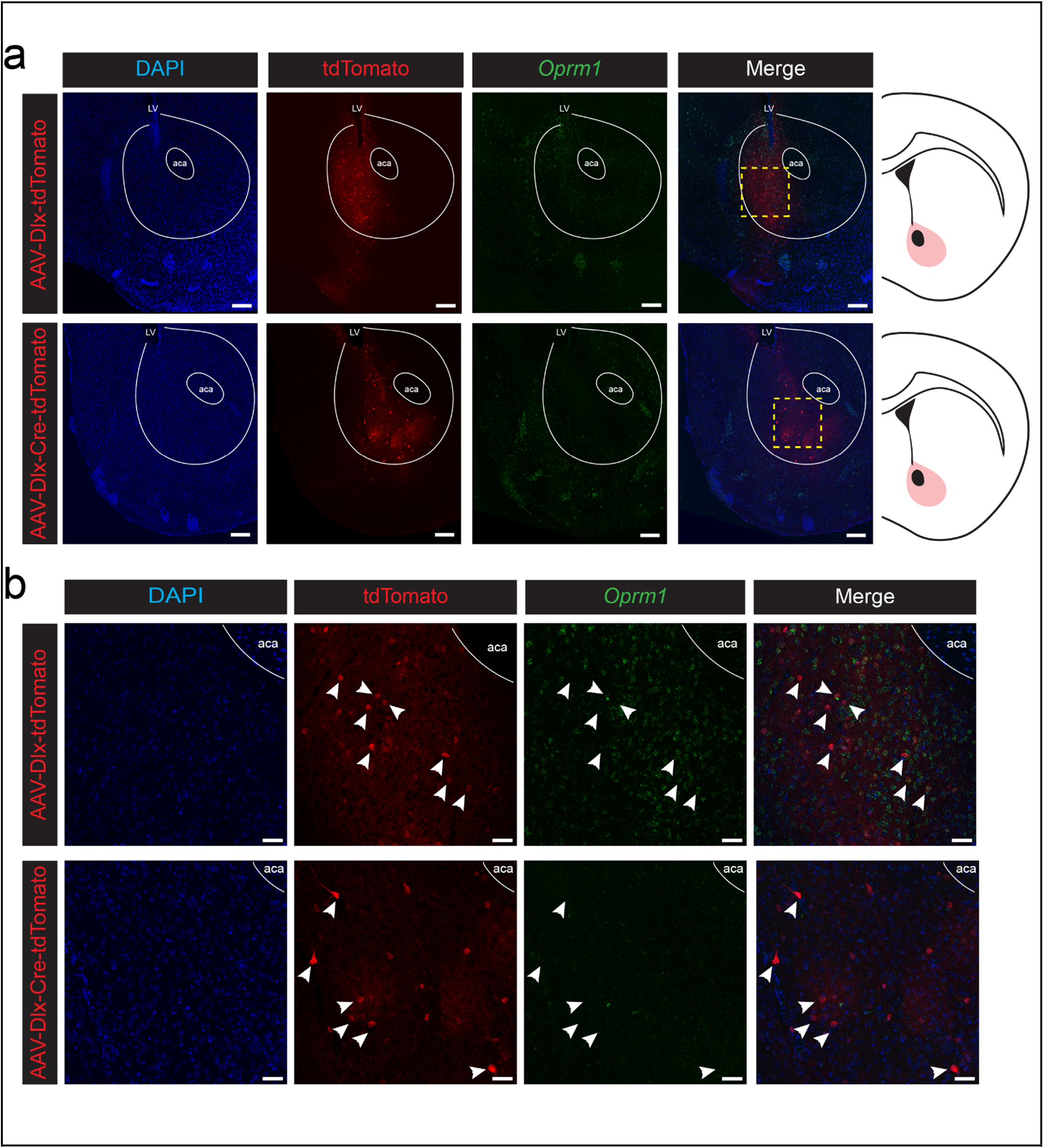
FISH validation of Oprm1 knockout from NAc interneurons using Dlx-iCre-tdTomato. (a) Zoomed-out representative images showing the entire right NAc from mice injected with either Dlx-tdTomato (top) or Dlx-iCre-tdTomato (bottom). DAPI nuclear stain in blue, endogenous tdTomato expression in red, *Oprm1* RNA in green, and the overlay. Scale bar= 200um (b) zoomed-in images of the yellow dashed squares in (a). Scale bar= 50 um. LV= lateral ventricle, aca= anterior commissure.

**Extended Data Figure 5.**
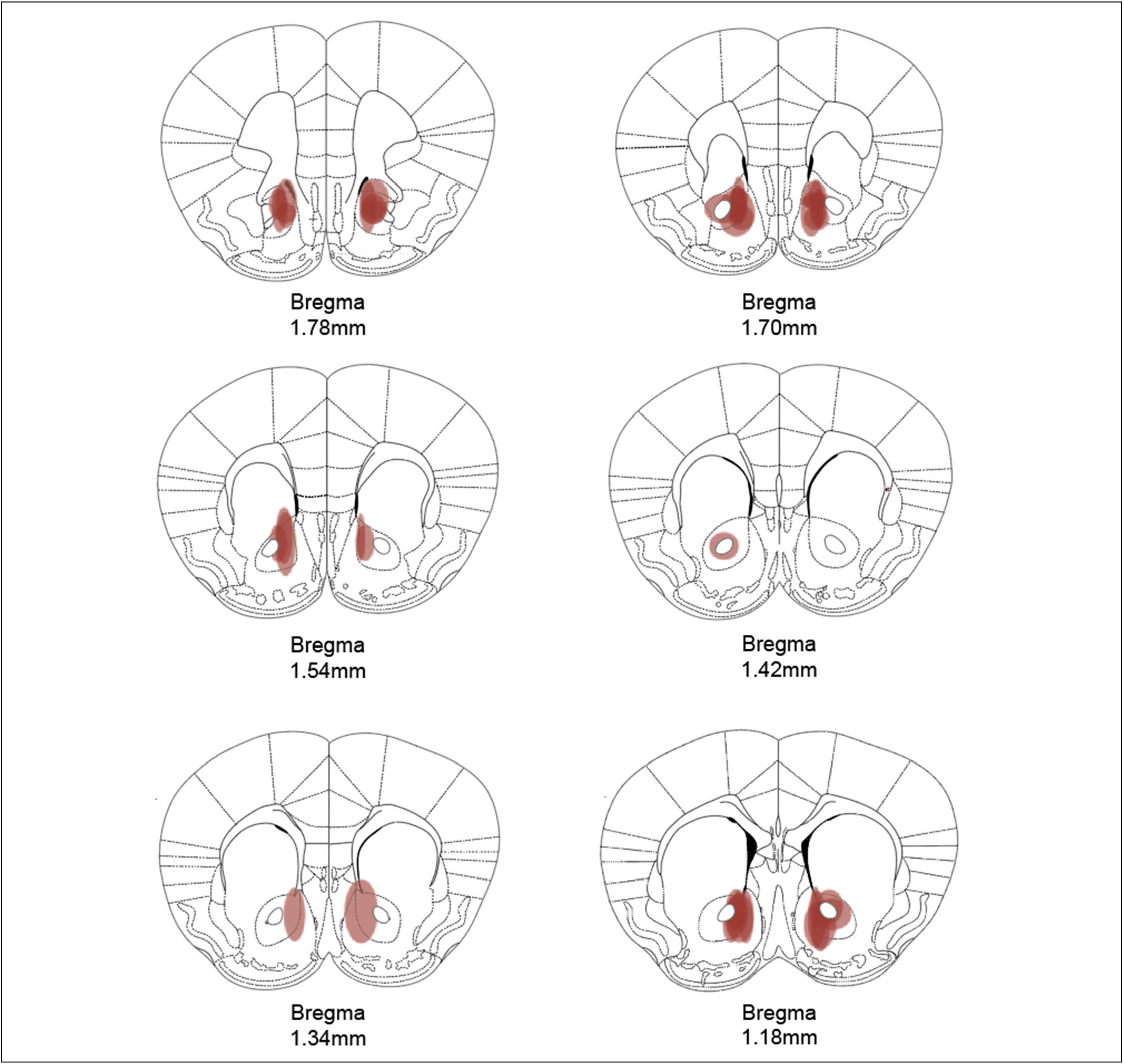
Viral expression patterns following nucleus accumbens injection of AAV8-mDlx-iCre-tdTomato.

**Extended Data Figure 6.**
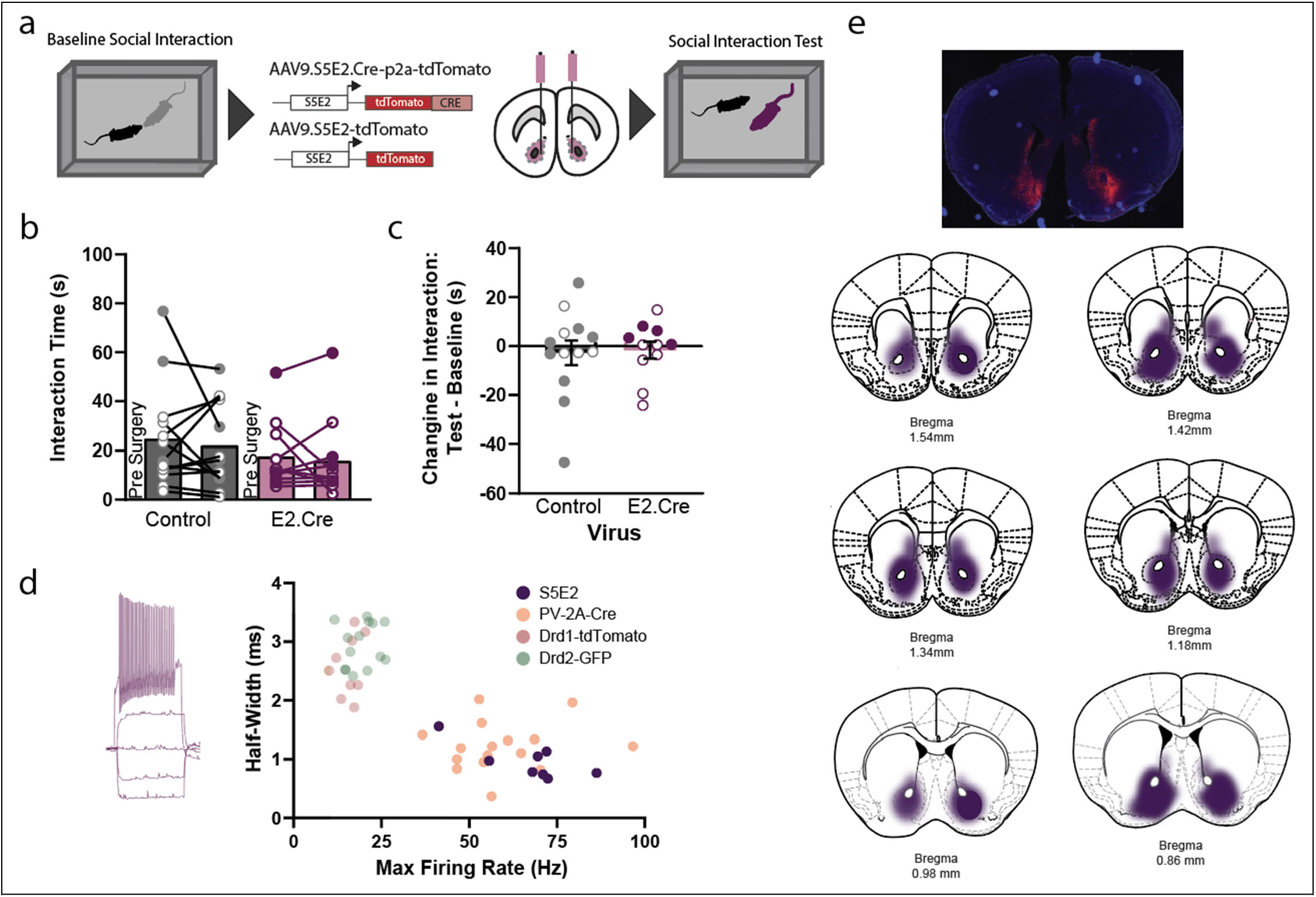
Conditional genetic knockout of Oprm1 from nucleus accumbens fast-spiking interneurons does not affect social interaction. (a) Illustration of experimental timeline: baseline social interaction tested before bilateral expression of Cre recombinase in the nucleus accumbens, followed by a social interaction test four weeks later. (b) Total interaction time before and after expression of Cre recombinase. (c) Change in social interaction expressed as a difference score between test and baseline session (n=11-13). (d) Example of whole-cell current-clamp recording from fluorescent neuron with fast-spiking properties (left), with electrophysiological properties of all cells with fluorescence driven by S5E2 enhancer sequence (n=8, right). Data from S5E2 cells are overlaid on published data^24^ showing electrophysiological properties of fast-spiking interneurons (labeled using PV-2A-Cre mouse line) and medium spiny neurons (labeled using Drd1-tdTomato and Drd2-eGFP reporter mice). (e) Representative image (top) and viral expression patterns in different coronal planes (bottom) following nucleus accumbens injection of AAV9-S5E2-Cre-P2A-tdTomato. Data are mean +/-SEM; open and filled circles represent female and male mice, respectively.

**Extended Data Figure 7.**
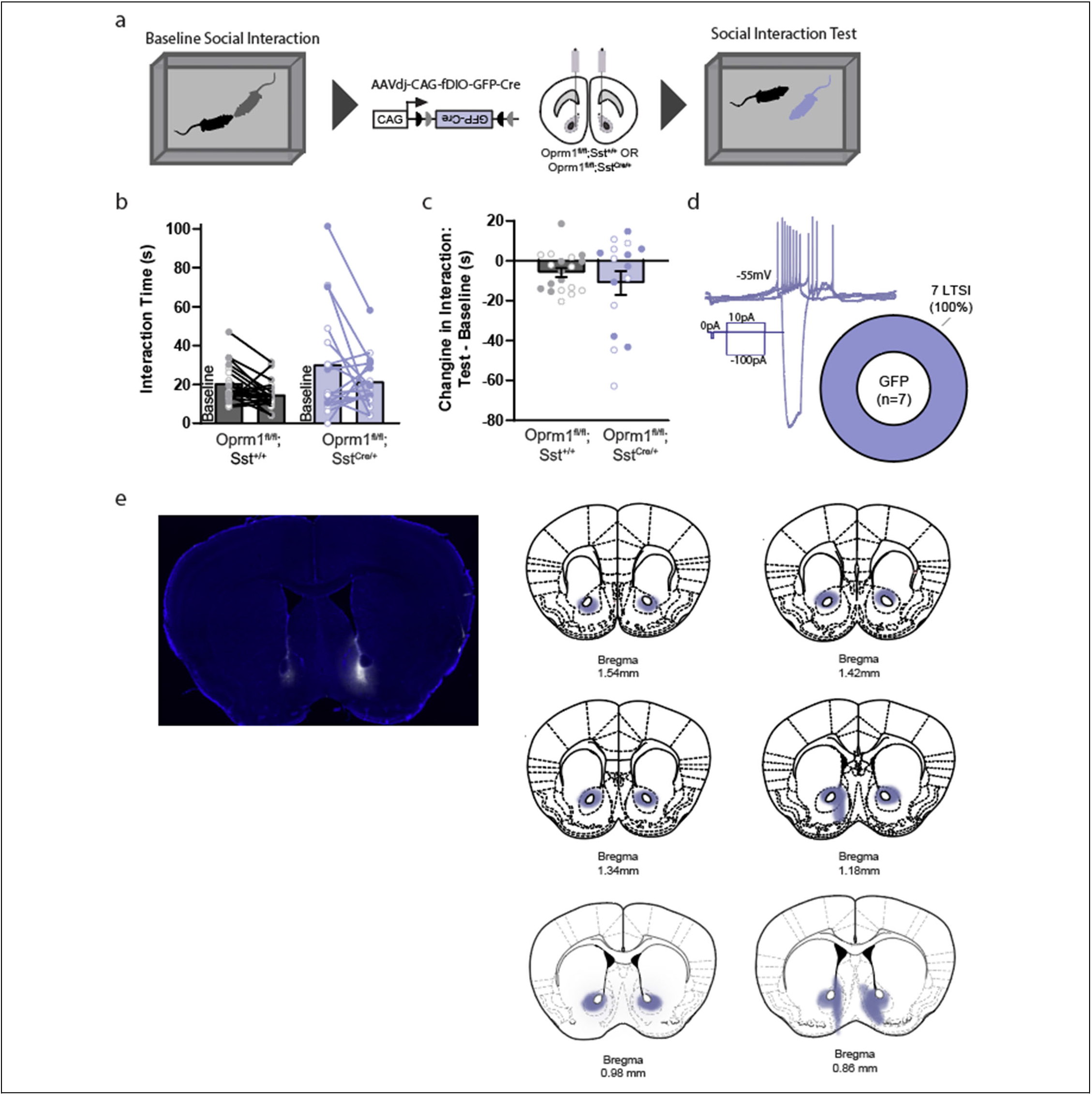
Conditional genetic knockout of Oprm1 from nucleus accumbens low threshold-spiking interneurons does not affect social interaction. (a) Illustration of experimental timeline: baseline social interaction tested before bilateral expression of Cre recombinase in the nucleus accumbens, followed by a social interaction test four weeks later. (b) Total interaction time before and after expression of Cre recombinase. (c) Change in social interaction expressed as a difference score between test and baseline session (n=16-19). (d) Example of whole-cell current-clamp recording from fluorescent neuron with low threshold-spiking properties, which were observed in 100% of green cells recorded (n=7). (e) Representative image (left) and viral expression patterns in different coronal planes (right) following nucleus accumbens injection of AAVdj-CAG-fDIO-GFP-Cre. Data are mean +/-SEM; open and filled circles represent female and male mice, respectively.

**Extended Data Figure 8.**
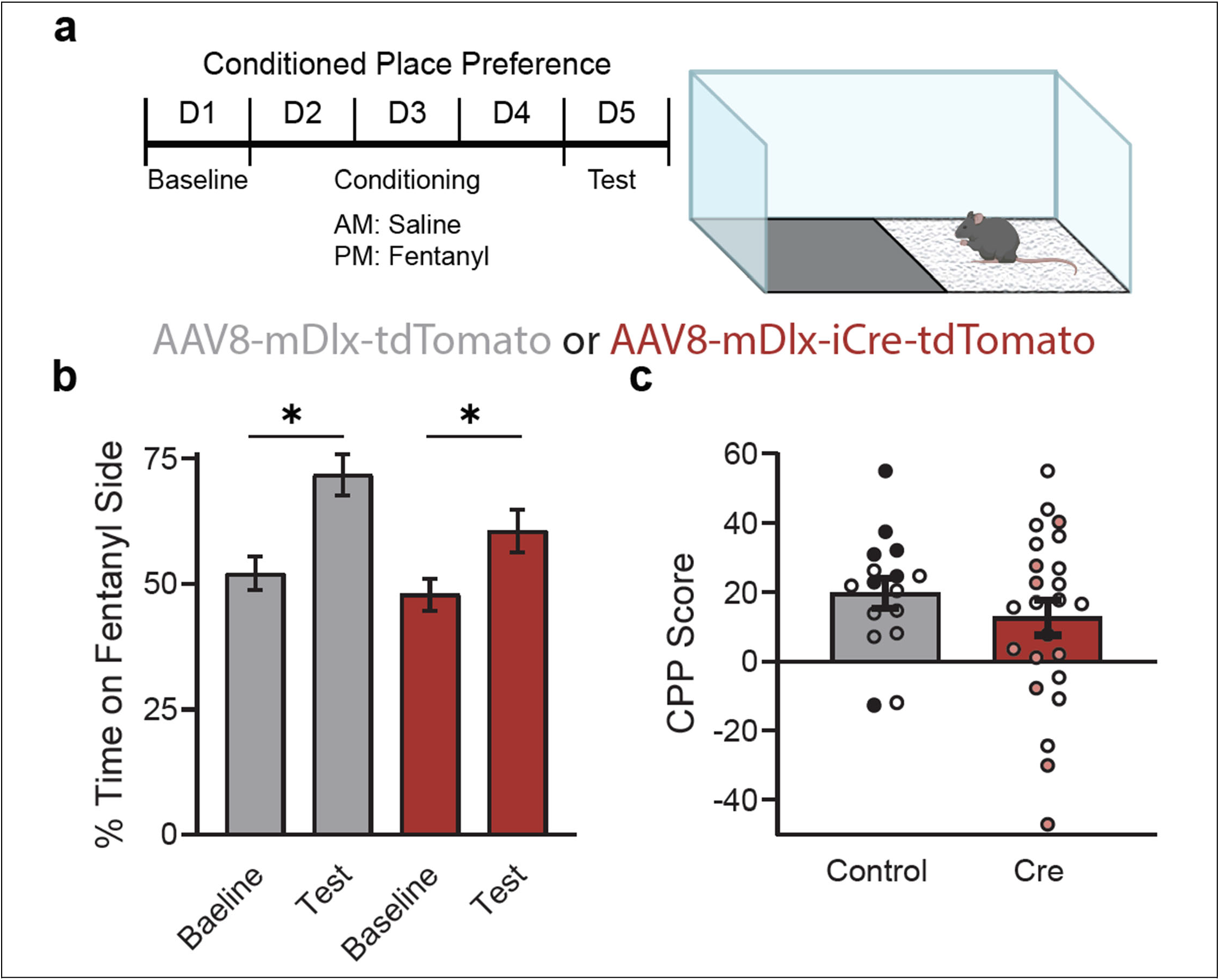
Conditional genetic knockout of Oprm1 from nucleus accumbens inhibitory interneurons does not affect fentanyl-conditioned place preference (CPP). Schematic of unbiased place conditioning procedure (a), with percent time on fentanyl side (b) and CPP score (c) after three conditioning sessions with fentanyl (0.04 mg/kg, s.c.). *p<0.05, ANOVA simple effect test for CPP within each group. Data are mean +/-SEM (n = 16-24); open and filled circles represent female and male mice, respectively.

